# Efficient Calculation of the Genomic Relationship Matrix

**DOI:** 10.1101/2020.01.12.903146

**Authors:** Martin Schlather

## Abstract

Since the calculation of a genomic relationship matrix needs a large number of arithmetic operations, fast implementations are of interest. Our fastest algorithm is more accurate and 25× faster than a AVX double precision floating-point implementation.

## 1 Background

The genomic relationship matrix (GRM) is the covariance matrix calculated from the SNP information of the individuals, i.e., from the minor allele counts [1]. It is an important ingredient in mixed models and generalized mixed models for analyses and predictions in genetics [2]. Let *n* be the number of individuals and *s* the number of SNPs per individual. Then, the calculation of the GRM needs of order *sn*^2^ arithmetic operations. Most software packages use floating-point arithmetics, for instance the `R` packages `AGHmatrix` [3], `qgg` [4], `rrBLUP` [5], `snpReady` [6], and `GENESIS` [7]. The software `GCTA` [8] treats missings explicitly. The package `SNPRelate` [9] also uses floating-point arithmetics for the covariance matrix, but uses bit manipulation algorithms for other calculations (identity-by-descent estimates). `PLINK` [10] profits from bit manipulations for calculating the uncentred covariance matrix.

In this paper we present a couple of ideas, how the GRM can be calculated efficiently from a SNP matrix. We assume that no values are missing. The emphasis will be on algorithms that allow for a vectorized implementation (SIMD) and that take into account that the entries of the SNP matrix are most efficiently coded by 2 bits, namely for the values 0, 1 and 2, as they are present in diploid organism under the common assumption of biallelic markers.

## 2 Results

The standard mathematical formula for the GRM requires floating point arithmetics. An algebraic reformulation shows that the cost intensive part involves only integers. Since the most elementary numbers need only a minimum of 2 bits, a diversity of approaches for the integer arithmetics is thinkable. The investigated methods are in brief:

- `Multiply`: uses 16-bit arithmetics
- `Hamming2`: uses pop counts (the number of bits that are 1)
- `ThreeBit`: uses a 3-bit representation and a single large hash table
- `TwoBit`: uses the 2-bit representation and two large hash tables
- `Shuffle`: similar to `TwoBit`, but with two tiny hash tables
- `Packed`: uses 4-bit arithmetics

They are all available through `crossprodx` in the package `miraculix` [11]. Tables 1 and 2 show that `Shuffle` is the fastest method, which is 35× faster than `crossprod` of `R` [12] in the SSSE3 implementation.

**Table 1:**
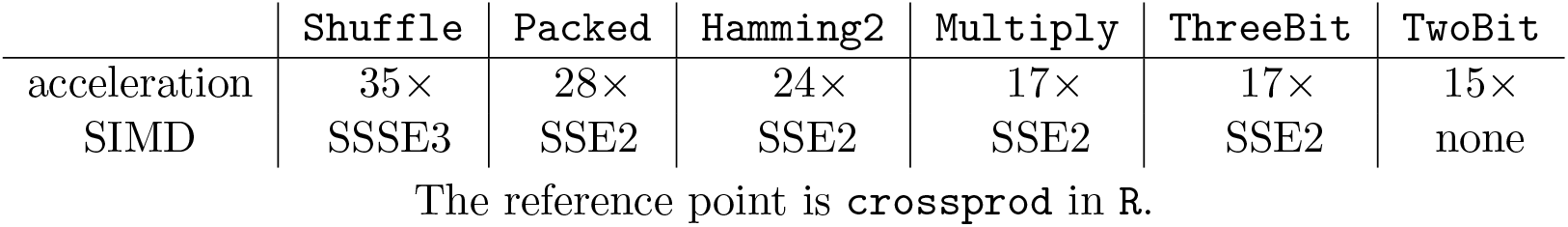
Accelaration of the calculation of *M*^T^*M* by `crossprodx`.

**Table 2:** Accelaration of the calculation of *M*^T^*M* by AVX2 implementations in `crossprodx`.

|  | **Shuffle** | **Packed** | **Multiply** | AVX (double) | AVX2 (32-bit integer) |
| --- | --- | --- | --- | --- | --- |
| acceleration | $48\times$ | $36\times$ | $29\times$ | $1.8\times$ | $4\times$ |

The reference point is **crossprod** in R.

The command `relationshipMatrix` in `miraculix` [11] for calculating the GRM is only negligibly slower than `crossprodx`. The `AVX2` variant of `Shuffle` is even 48× faster than `crossprod` [12] and 48*/*1.8 *≈* 25× faster than a standard AVX double precision implementation for calculating the crossproduct of an arbitrary matrix, cf. Table 2. Furthermore, our algorithms have not even any cummulative rounding error.

Tables 1 and 2 also show that the `AVX2` performance is hard to predict from the SSE performance. AVX2 variants for `TwoBit` and `ThreeBit` are not given since full vectorization is not possible. `Hamming2` has not been persued because of its memory demand.

For the benchmarks, we used an *s* × *n* SNP matrix with *n* = 1000 individuals and *s* = 5 *·* 10^5^ SNPS. The calculations were performed on an Intel(R) Core(TM) i7-8550U CPU @ 1.80GHz with R version 3.6.0 on Xubuntu. Although the code in `miraculix` is parallelized, we used only a single core for the benchmarks. Nonetheless, the AVX2 variant of `Shuffle` takes not more than 7 seconds. The code for the benchmarks is available from the man page of `crossprodx` in `miraculix`.

## 3 Discussion

First, with respect to the memory needs of the SNP matrix, algorithms that use the 2-bit representation of a SNP value should be preferred. Among them, we have a sequence of distinct algorithms that differ in their speed-up and their SIMD requirements: `TwoBit` (15×; SIMD not used), `Packed` (28×; SSE2); `Shuffle` (35×; SSSE3).

Second, the use of perfect hash tables to cut calculations short might be of general importance. Third, since loading from non-aligned memory allocation is reported to be slower [13], the package miraculix was designed to avoid non-aligned loadings. Tests on the implemented package however show that the running time by non-aligned loadings is not reduced for SSE implementations. The speed is reduced by 5 to 10 % in AVX2 implementations. As the compressed SNP matrix is made available to the user as an R object and as the memory allocation by R is only 32-bit aligned, additional memory is allocated and the SNP matrix is aligned to 128 or 256 bits. Furthermore, additional zeros are appended so that the virtual number of SNPs is a multiple of the number of the SNPs that can be treated in a single step. The storing formats of `TwoBit`, `Multiply`, `Packed`, and `Shuffle`, including their AVX2 variants, were made compatible, i.e., the allocations are all based on a 256-bit alignment. A check and a reallocation are implemented for the case that memory is moved. This might happen when the garbage collector `gc` is called by `R`, for instance.

## 4 Conclusion

The combination of algebraic reformulation, bit manipulations and hash tables can reduce largely the computing time on SNP data. In the case of calculating the GRM, the computing time could be reduced by factor 25 in comparison to a straightforward AVX double precision implementation. As a spectrum of implementations exist, there is a chance of further improvement and of further applications of the underlying ideas.

## 5 Methods

Let *M* be an *s × n* SNP matrix of *n* individuals and *s* SNPs. We need to consider only the fast calculation of the crossproduct *M*^T^*M*, since the GRM *A* can be calculated from *M*^T^*M* at low costs. This can be seen as follows.

Let **1**_*k*_ be the vector of length *k* whose components are all equal to 1. The centred and normalized GRM *A* is calculated as

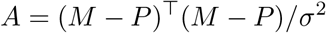

where

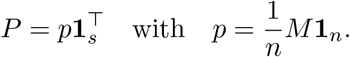

and

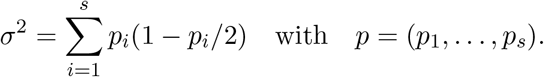

Note that replacing the value *p*_*i*_ by the allele frequency 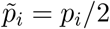, we have the usual formula for *σ*^2^,

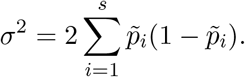

Let *B* = *M*^T^ *M*1_n_. Then

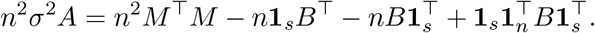

Hence, the integer-valued matrix *n*^2^*σ*^2^*A* can easily be calculated from the matrix *M*^T^*M* without any numerical error and at low computational costs of order *n*^2^. Now,

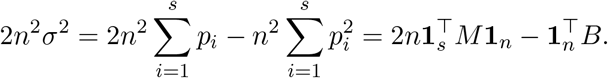

Again, 2*n*^2^*σ*^2^ can easily be calculated from *M* and *M*^T^*M* without any numerical error. The computational costs are of order *n*(*n* +*p*), hence still some magnitudes smaller than the costs of calculating the crossproduct *M*^T^*M*.

### 5.1 Algorithms for scalar products

Instead of considering the crossproduct *M*^T^*M*, it suffices to consider the scalar product of two vectors *a* = (*a*_1_*, …, a_s_*) and *b* = (*b*_1_, …, *b*_*s*_) whose components *a*_*i*_ and *b*_*i*_ have the values 0, 1 or 2. For simplicity and clarity, we will primarily refer to SSE commands in the following, and not to AVX.

#### 5.1.1 Simple Multiplication

An immediate way of calculating the scalar product from a compressed 2-bit representation is to extract the first two bits of each of the two vectors *a* and *b* and to continue with integer arithmetics. Then the next two bits are extracted using shifting, and so on. Clearly, this procedure can be vectorized. Of particular advantage here is the SSE2 command `_mm_madd_epi16`, which multiplies and adds two consecutive 16-bit integers so that only 7 shifts are necessary for a vector of 64 SNP values, i.e., for 128 bits. We call this method `Multiply`.

#### 5.1.2 Hamming Distance

The algorithm used in `PLINK` [14, 10] is based on the idea that a value is represented by the number of bits that equal 1 in a 4-bit representation. The values of the vectors *a* and *b* must be coded asymmetrically by two mappings *f*_*a*_ and *f*_*b*_, say, as a coding by single mapping is not possible. Then, the bitwise &-operator is applied before the number of 1’s is counted. Table 3 gives a possible realisation.

**Table 3:**
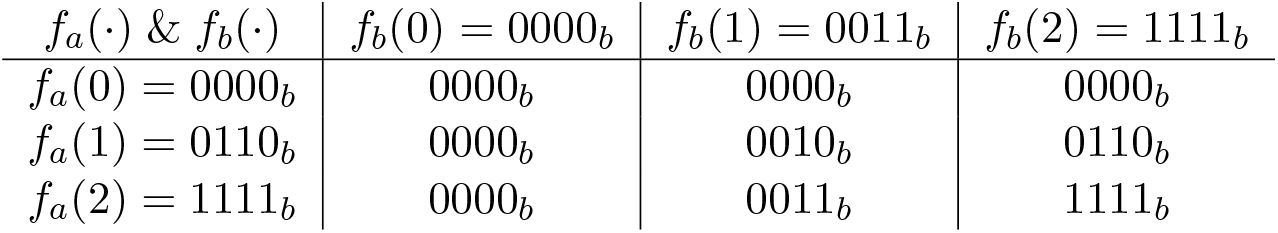
Table of values for the Hamming distance method.

| $f_a(\cdot) \& f_b(\cdot)$ | $f_b(0) = 0000_b$ | $f_b(1) = 0011_b$ | $f_b(2) = 1111_b$ |
| --- | --- | --- | --- |
| $f_a(0) = 0000_b$ | 0000<sub>b</sub> | 0000<sub>b</sub> | 0000<sub>b</sub> |
| $f_a(1) = 0110_b$ | 0000<sub>b</sub> | 0010<sub>b</sub> | 0110<sub>b</sub> |
| $f_a(2) = 1111_b$ | 0000<sub>b</sub> | 0011<sub>b</sub> | 1111<sub>b</sub> |

The number of bits that equal 1 can be calculated by SSE2 commands based on work by [15, 16]. We call the method `Hamming2`. (An SSSE3 implementation in `miraculix` [11] is called `Hamming3`.) The method can be turned into a particularly fast implementation when novel AVX512 commands are used for the pop counts, e.g. `_mm512_popcnt_epi64`. See also `SNPRelate` [9] for pop count implementations. Still, the storing costs of the SNP matrix *M* remain high, namely 2 × 4 = 8 bits per SNP in a standard implementation.

#### 5.1.3 Perfect Hash Table

Let us consider the product of the first elements *a*_1_ and *b*_1_ of the two SNP vectors *a* and *b*. Let us code the SNP values 0, 1 and 2 by 3 bits, e.g. as 000_*b*_, 011_*b*_ and 110_*b*_, respectively, and denote this mapping by *f*. Then, a perfect hash table for *f* (*a*_1_) & *f* (*b*_1_) returns 0, 1, 2, 4 for 000_*b*_, 011_*b*_, 010_*b*_, and 110_*b*_, respectively, cf. Table 4.

**Table 4:** Table of values for the `ThreeBit` method.

| $f(\cdot) \& f(\cdot)$ | $f(0) = 000_b$ | $f(1) = 011_b$ | $f(2) = 110_b$ |
| --- | --- | --- | --- |
| $f(0) = 000_b$ | 000<sub>b</sub> | 000<sub>b</sub> | 000<sub>b</sub> |
| $f(1) = 011_b$ | 000<sub>b</sub> | 011<sub>b</sub> | 010<sub>b</sub> |
| $f(2) = 110_b$ | 000<sub>b</sub> | 010<sub>b</sub> | 110<sub>b</sub> |

Subvectors of length *k* of *a* and *b* can be treated in the same way and the perfect hash table returns then the scalar product of the subvectors. The hash table will be indexed by 3*k*-bit numbers, i.e. by values between 0 and 2^3*k*^ − 1. Since *k* should be as large as possible at a smallish size of the hash table, and 3*k* bits should fit nicely into 1, 2 or 4 bytes, the only reasonable choice for *k* is *k* = 5, so that 15 bits in a 16-bit representation of a vector with *k* = 5 components are used. The precise size of the hash table is then 110 110 110 110 110_*b*_ + 1 = 28087 bytes. We call this method `ThreeBit`.

#### 5.1.4 Two Perfect Hash Tables

Since we did not find a simple way to use a single hash table based on a 2-bit representation of the SNP values, we consider here two hash tables and the two bitwise operators & and *|*. The first hash table should return 1 and 4 for 01_*b*_ and 10_*b*_, respectively, while the second hash table returns 2 for 11_*b*_. All other values in the hash tables are 0, cf. Table 5.

**Table 5:**
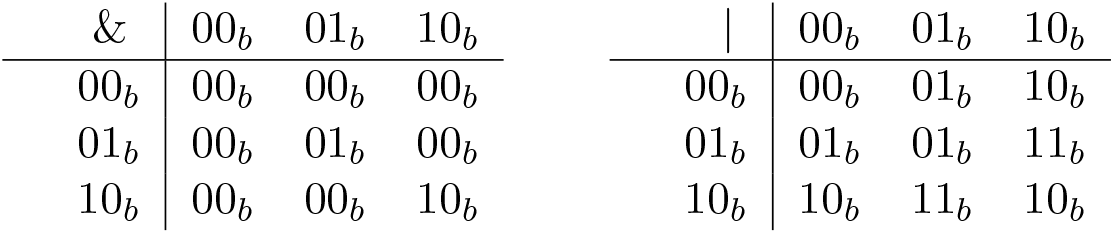
Tables of values for the `TwoBit` method.

Then, the sum of the two table values yields the product *a*_1_*b*_1_. Scalar products of subvectors of length *k* can also be treated by two hash tables. Since the size of both hash tables is of order 2^2*k*^, one possible choice is *k* = 8, so that the size of the second hash table is 65536 bytes. We call this method `TwoBit`.

The disadvantage of `TwoBit` (and `ThreeBit`) is that the look-up in the hash table prohibits a full vectorization. A much better choice is therefore *k* = 2: the SSSE3 command `_mm_shuffle_epi8` looks 16 values up at once in a hash table of size 16. We call this variant `Shuffle`.

#### 5.1.5 Packed arithmetics

A last idea is to emulate a multiplication by bitwise operations and partial sums. Let ≫ denote the bitwise shift operator and let

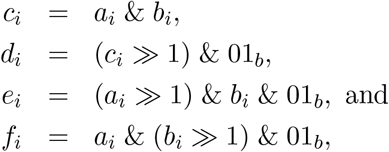

so that *c*_*i*_ + 2*d*_*i*_ = *a*_*i*_*b*_*i*_ if *a*_*i*_ = *b*_*i*_ and 0 else. Furthermore 2(*e*_*i*_ + *f_i_*) = *a*_*i*_*b*_*i*_ if *a*_*i*_*b*_*i*_ = 2 and 0 else. In total, we have *c*_*i*_ + 2*d*_*i*_ + 2(*e*_*i*_ + *f_i_*) = *a_i_b_i_*. Let *g*_*i*_ = *d*_*i*_ + *e*_*i*_ + *f_i_*. Since *g*_*i*_ = *d*_*i*_ | *e*_*i*_ | *f*_*i*_ for the bitwise operator *|*, only the values of *g*_*i*_ and *c*_*i*_ need to be summed up. An immediate extraction of the values of *g* = (*g*_1_*, …, g_s_*) and *c* = (*c*_1_*, …, c_s_*) by shifting as in the `Multiply` algorithm would be rather expensive. Instead, a 4-bit arithmetic can be introduced in an intermediate step for the four vectors

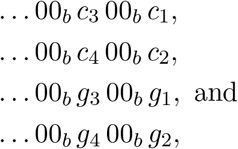

which can be obtained by two shifts and 4 bitwise &-operations in total, if the ordering in the memory is *… c*_4_ *c*_3_ *c*_2_ *c*_1_. Since the value of *c*_*i*_ is at most 2, a sevenfold summation of the first two displayed vectors leaves each component within its 4 bits (using any unsigned integer SIMD addition). Since the value of *g*_*i*_ is either 0 or 1, even a fifteen-fold summation is possible for the last two displayed vectors. Afterwards, the 4-bit values are extracted and further summed up. We call this method `Packed`.

The novel AVX512 command `_mm512_popcnt_epi64` might improve this approach, as it allows to count the number of bits being 1 in *c_i_*, *d*_*i*_ and *e_i_ | f_i_*, so that the number of products is counted that equal (i) 1^2^ or 2^2^, (ii) 2^2^, and (iii) 1 *·* 2 or 1 *·* 2. This variant can be seen as a 2-bit analogue of the `Hamming2` algorithm, and will be implemented in future.

## 6 Abbreviations

AVX: Advanced vector extensions
SIMD: Single instruction, multiple data
SNP: Single nucleotide polymorphism
SSE: Streaming SIMD extensions
SSSE: Supplemental streaming SIMD extensions
GRM: Genomic relationship matrix

## 7 Acknowledgements

The author is grateful to Torsten Pook, Christopher Dörr and Alexander Freudenberg for hints and for comments.

## Notes

https://github.com/schlather

